# Biophysical characterisation of the structure of a SARS-CoV-2 self-amplifying - RNA (saRNA) vaccine

**DOI:** 10.1101/2022.10.03.507132

**Authors:** Daniel P Myatt, Lewis Wharram, Charlotte Graham, John Liddell, Harvey Branton, Claire Pizzey, Nathan Cowieson, Robert Rambo, Robin J Shattock

## Abstract

The current SARS-Covid-2 pandemic has led to an acceleration of messenger – ribonucleic acid (mRNA) vaccine technology. The development of production processes for these large mRNA molecules, especially self-amplifying mRNA (saRNA) has required concomitant development of analytical characterisation techniques. Characterising the purity, shape and structure of these biomolecules is key to their successful performance as drug products. This paper describes the biophysical characterisation of the Imperial College London Self-amplifying viral RNA vaccine (IMP-1) developed for SARS-CoV-2. A variety of analytical techniques have been used to characterise the IMP-1 RNA molecule. In this paper we use UV spectroscopy, dynamic light scattering (DLS), size-exclusion chromatography small angle scattering (SEC-SAXS) and circular dichroism (CD) to determine key biophysical attributes of IMP-1. Each technique provides important information about the concentration, size, shape, structure and purity of the molecule.

**Statement of significance:** This paper is highly significant as it provides a prescient biophysical characterisation of an efficacious Sars-Cov-2 vaccine self-amplifying (sa)RNA molecule. RNA vaccines have been a major scientific breakthrough of the Covid-19 pandemic. saRNA is a further development of conventional mRNA vaccines, amplifying the RNA of interest in the cell, allowing the vaccine to be administered at lower dosages. These new biologics are distinct from previous biologics and have required distinct analytical characterisation. The analytics described herein provide detailed information on the size, shape, and structure of the RNA molecule. This paper is therefore an important step in characterising large saRNA biological relevant molecules.

## 1) INTRODUCTION

The SARS-Cov-2 pandemic has caused untold disruption worldwide and has claimed a current global death toll of several million people (1). Several SARS-Cov-2 vaccines, including mRNA vaccines, are now in development, in clinical trials, at approval stage or approved (2). One major scientific advancement from the pandemic has been the development of mRNA vaccine technology. These vaccines work by supplying an mRNA copy of the target antigen, typically the SARS-Covid 2 spike protein, to the host cell translation machinery (3). Self-amplifying RNA vaccines are a step forward in RNA vaccine development. They work by adding a self-amplifying (sa) RNA replicon to the RNA molecule, causing the host cell to multiple the copy number of the target antigen RNA (4). This *in vivo* amplification allows the saRNA vaccine to be delivered at significantly lower doses, typically 10-100 fold less in comparison to conventional mRNA vaccines. However, the extra self-amplifying code means the RNA molecules are significantly larger than conventional mRNA vaccines at around 9-12 kbps (5). In this paper, we analyse a saRNA vaccine developed by the Imperial College London (ICL) team, we call IMP-1. It consists of the genetic code for a Venezuelan equine encephalitis virus (VEEV) replicase in addition to a pre-fusion stabilized spike protein of SARS-CoV-2 (6). This molecule has proven efficacious in pre-clinical mouse studies and immunogenic in initial clinical trials (Clinical trial: ISRCTN17072692 (II)) (6).This vaccine was manufactured at scale by the Centre for Process Innovation (CPI) in a program to develop, manufacture and supply the saRNA COVID-19 vaccine developed by ICL, with funding provided by the Department of Business, Energy and Industrial Strategy (BEIS) via the UK Vaccine Taskforce (VTF) (7).

The scale up of the IMP-1 involved CPI developing a variety of analytical methods to characterise the IMP-1 molecule. One great strength of RNA vaccines is the ability to quickly design and manufacture the test molecule. This allowed the IMP-1 RNA molecule to be developed for pre-clinical testing, within 14 days of the release of SARS-CoV-2 genetic data (8). RNA vaccines can also be altered quickly to counter new variants and are an ideal platform to reduce the time for development, manufacture and testing of vaccines to within a 100-day goal (9). The mRNA vaccine production platform is being standardised to incorporate several steps. Firstly, the RNA molecule is synthesised in an *in vitro* transcription (IVT) cell free synthesis by the addition of the DNA plasmid of interest and a cocktail of reagents including an RNA polymerase, ribonucleotide triphosphates (rNTPs), RNase inhibitors, MgCl_2_, a pyrophosphatase, a pH buffer containing an antioxidant, and a polyamine at the optimal concentrations, plus a capping analogue if co-translational capping is required in this or a further separate step (10, 11). This reaction is optimised for reagent concentrations, time and temperature. After the RNA molecule is synthesised a purification and polishing step is performed to remove unused NTP’s, enzymes and DNA fragments. This step has classically used a salt or solvent precipitation step at laboratory scale (12), but chromatographic capture and release is becoming the method of choice for scale up (13). The RNA product can then be exchanged into the required buffer and concentrated using diafiltration/ultrafiltration (DF/UF), followed by encapsulation in a lipid nanoparticle (14). The active drug product is then stored ready for transport. Each step in this process contains the potential to generate incomplete drug substance, through incomplete transcription, hydrolysis and degradation of the RNA molecule. Analytical quality controls of the RNA molecule are therefore critical to allow the identification of alterations to the molecule and allow mitigation and prevention strategies to be put into place. A variety of analytical methods have been used to characterise the RNA drug substance in process including visual appearance, subvisible particles, UV and fluorescence spectroscopy, agarose gel electrophoresis, capillary electrophoresis (CE), chromatography (SEC – size-exchange chromatography, reverse-phase chromatography (RP), anion-exchange chromatography (AEX)), light scattering (dynamic light scattering – DLS, SLS – static light scattering, MALS – multi-angle light scattering), circular dichroism (CD), reverse transcriptase PCR (RT-PCR) and liquid chromatography - mass spectrometry (LS-MS) (15, 16).

RNA structure differs from DNA (deoxyribonucleic acid) in three fundamental ways. Firstly, its backbone contains the ribose sugar. Ribose differs from deoxyribose, in that it has an addition hydroxyl in the 2’-position. Secondly, RNA uses uracil, instead of thymine as one of its bases. Uracil lacks a methyl group in the 5’-position, in comparison to thymine. Finally, RNA is usually single stranded (17). These differences mean RNA exists in very different structures in comparison to DNA. The single stranded (ss) RNA forms secondary structure such as double helices by complementary base pairing. In addition to Watson-Crick base pairing other non-Watson-Crick base pairing, such as G:U Wobble base pairing, are possible (18). Due to the steric hinderance of the 2’-hydroxyl, RNA double helix more closely resembles DNA A-form helix. To maximise these complementary regions various stem loop structures, such as hairpins, bulges or simple loops are possible. RNA has enormous rotational freedom in the backbone of its non-base paired regions allowing it to form complex tertiary structures such as helical duplexes, triple-stranded structures and other structures (19). A variety of biophysical techniques, including NMR (nuclear magnetic resonance) (20), EM (electron microscopy) (21), X-ray-crystallography, and SAS (small angle scattering) (22) have been used to determine RNA structures to varying resolution. Many of the known RNA structural motifs have been determined from the limited number of atomic resolution structures available (23). In recent years biophysical data linked with biochemical data such as chemical probing, sequencing (24), computational modelling (25) and predication methods (26) have further aided our understanding of RNA structure. Whilst these methods have proved successful for short length RNA, longer RNA, such as saRNA, have increasingly complex structures, that will need extensive experimental validation. RNA structure is also highly anionic due to the phosphate backbone (pKa ≈ 2). The structure is stabilised by the addition of cations, most commonly the divalent cation magnesium (Mg_2+_) (27). The exact effect of salt on the RNA is RNA and salt concentration dependant. At dilute concentrations, cations allow RNA to form more compact structures and so decrease its size (28–31). At higher concentrations, as used in RNA manufacture, the addition of salt causes intramolecular RNA base pairing leading to aggregation (28).

RNA has been extensively characterised using UV spectroscopy to determine the concentration and purity of solutions. The absorbance spectra and extinction co-efficient (ε) for each individual RNA nucleotide has been determined. If no experimentally determined extinction-co-efficient at 260 nm (ε260) is available, theoretical calculations are available. General conversion factors for single stranded (ss)RNA of 40 µg/mL per 1 A(absorbance) unit are often used, although for oligonucleotides more accurate A_260_ extinction co-efficients (ε_260_) can be determined using base composition (bc) and nearest neighbour (nn) sequence specific calculations. Using Beer-Lambert with an accurate extinction-co-efficient at 260 nm (ε260) an RNA concentration in solution can be determined quickly. Determining the ratio between absorbances at 260 and 280 nm (A_260_ : A_280_) is useful in determine RNA purity, particularly for protein contaminants. A ratio equal or greater than 2 is indicative of pure RNA, when the assay is performed in mildly alkaline (pH = 8 – 8.5), low ionic strength buffers (32).

Dynamic Light Scattering (DLS) is a solution based, non-invasive scattering technique for determining the size and polydispersity of particles in a solution. DLS involves measuring the Brownian motion of particle in solution via light scattering fluctuations. Analysis of these particle fluctuations yield the translation diffusion co-efficient (D), a measure of the velocity of the Brownian motion of the particles in the sample. Using the Stokes-Einstein equation the translation diffusion co-efficient (D) can be converted to the particle size. The sample particle size is commonly given as the Z-Average diameter (Z_D_) defined as the intensity-weighted mean hydrodynamic diameter. With the hydrodynamic diameter (D_h_) being defined as the diameter of a hard sphere that diffuses at the same speed as the particle or molecule being measured (33). Alongside Z_D_, the polydispersity index (PDI) of the sample is given. PDI is a dimensionless measure of the broadness of the distribution of the particle sizes derived from cumulant analysis of the data. PDI values range from 0 to 1, with higher values indicative of greater polydispersity.

Biological small angle x-ray scattering (BioSAXS) experiments provide information on the size and shape of biological sample molecules such as RNA (34). They are high throughput experiments that can be performed either at off-line home X-ray sources or more commonly at synchrotron facilities (35) with limited amounts of purified sample (34). In BioSAXS experiments, a highly collimated X-ray beam is fired at the sample of interest. A small number of the X-rays are scattered by collision with the atoms of the molecules in the sample. A detector collects the scattering data at low angles (0.1-10°). This 2-dimensional data is then radially averaged, and the buffer subtracted to give a one-dimensional plot of scattering intensity I(q) as a function of the scattering angle vector (q). As a scattering technique, BioSAXS is disproportionally affected by large aggregates, with the scattering signal being proportional to the square of the molecular weight (36). This means that preliminary characterisation techniques such as agarose gels, UV spectrometry (A _260:280_) and dynamic light scattering (DLS) are critical to confirm sample size and purity. BioSAXS can also be linked directly to a high-performance liquid chromatography (HPLC) system. This allows HPLC-SEC (high-performance liquid chromatography - size exclusion chromatography) separation of the biological sample based on hydrodynamic radius (R_H_) to be performed, before SAXS (37). The BioSAXS is then recorded in timed frames as the sample solution passes through the x-ray beam, and the buffer and sample frames can be grouped and averaged to give the 1D SAXS curve. BioSAXS, relies on the scattering contrast between the buffer, which is usually aqueous in nature (∼ 344 e/nm^3^), and the biomolecule of interest, such as RNA (550 e/nm^3^). HPLC-SEC allows the precise matching of the experimental buffer used in the buffer and RNA sample, without the need for sample dialysis, as both samples will be measured using the same HPLC running buffer.

BioSAXS data is analysed using several methods. The most common is these is the Guinier plot, developed by Andre Guinier in 1939 (38), which states: -

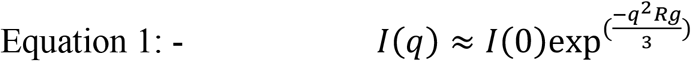

Where,

I(q) = intensity at the scattering vector q

I(0) = Intensity at zero angle

q = scattering vector angle

Rg = radius of gyration

The equation can be plotted as ln(I) versus q^2^, using data points in the Guinier region, which for globular molecules is given as q.Rg 0.5 (min) – 1.3 max. The plot intercept determines the intensity at zero angle I(0), which is proportional to the molecular weight of the particles and the slope determines the radius of gyration (R_g_), which provides an estimate of the molecule size as determined from the mean distance from the centre of mass weighed by the contrast of the electron density (39). Guinier plots at a range of sample concentrations are an important method in spotting molecule aggregation, interparticle interference and concentration dependant complexes (40). Other commonly used BioSAXS plots are the distance distribution function (PDDF) and Kratky. The PDDF plot is an inverse Fourier transform of the scattering plot. PDDF plots are useful in determine the general shape of the molecule, with characteristic shapes being easy to spot. It also determines key parameters such as radius of gyration (Rg) and maximum diameter (D_max_) of the molecules in the sample. Dimensionless Krakty (q.Rg^2^ x I(q)/I(0) versus q.Rg) plots give a semi-quantitative analysis of the flexibility of the molecule analysed. Compact molecules display parabolic bell-shaped shaped curves, whilst unfolded molecules show hyperbolic curves that plateau out at high-q values. Semi-compact molecules display a combination of the two curves (41).

Circular dichroism (CD) is a relatively quick, solution-based spectroscopic technique that requires few consumables and small amounts of purified (>95%) samples to obtain orthogonal experimental information of the structure and conformation of chiral biomolecules such as RNA (42). CD is calculated by subtracting the absorbance the absorbance of left versus right circular polarised light:-

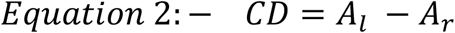

Where,

CD = circular dichroism

A_l_ = absorbance of left circular polarised light

A_r_ = absorbance of right circular polarised light

CD experiments follow Beer-Lambert and can be tailored to concentration and cell pathlength, with an optimal theoretical absorbance (A) of ∼0.869 (43). Several different types of experiments can be performed by CD, including sample batch-to-batch comparison (15), the effect of salts and solvent (44), buffers and pH (44), and temperature ramps (45) to name a few. Although care needs to be taken as to the UV transparency of additives, with sodium phosphates buffers being commonly used (42).

Deoxyribonucleic acid (DNA), has been extensively characterised by CD in the literature (46). There is much less literature on the circular dichroism of ribonucleic acid (RNA), particularly large saRNA molecules, such as IMP-1. The ultra-violet (UV) CD spectral region, from 170nm to 320nm, contain several of RNA absorption bands of interest. The positive band CD signal at approximately 265 nm is known to relate to intra- and intermolecular and base-stacking and base-pairing of the RNA molecule, including Watson–Crick and non-Watson-Crick base pairs (47). This positive band lies within a broader band from 210 to 300 nm, due to interactions between the RNA bases and the sugar phosphate backbone (48), (49). As expected, changes of the CD spectra in this region are known to relate to sequence differences affecting base pairing and stacking of the RNA molecule. The CD absorptions bands at the far-UV end of the spectrum are known to relate to the helical structure of the RNA molecule. A prominent positive band at ∼ 185 nm and a negative band at ∼ 205 nm is indicative of right-handed RNA, such as A-form RNA. Whilst a positive band at ∼ 180 nm and a negative band at ∼ 190nm is indicative for left-handed nucleic acids such as Z-form RNA (50).

## 2) MATERIALS AND METHODS

All chemicals and biologics purchased were of an analytical grade or higher.

### 2.1) *In vitro* transcription and purification of IMP-1

The IMP-1 mRNA sequence (6) was transcribed from a plasmid containing self-amplifying sequences from Venezuelan equine encephalitis virus (VEEV) and a Pre-fusion stabilized spike protein of SARS-CoV-2. The IVT reaction was performed as per McKay et al. (6) using plasmid DNA purchased from Aldevron, UK with the reaction optimised as per the design-of-experiment (DOE) methodology described by Samnuan (11) and the capping using the CleanCap ® Reagent AG performed as per the manufactures protocol (Trilink Biotechnologies, USA). After the IVT reaction was complete the DNA template was digested using DNase 1 enzyme (New England Biolabs, UK). The reaction mixture was purified, buffer exchanged and concentrated using Tangential Flow Filtration (TFF) and chromatography steps. The IMP-1 RNA in 5 mM Sodium Citrate, pH 6.8 buffer was stored at -80 °C.

### 2.2) A_260_ /A_280_ UV spectrometry assay

The concentration and purity of IMP-1 was determined by its A_260_ absorption value and the ratio of its A_260/280_. Fifty-fold diluted samples were measured in 10mM Tris buffer, pH 8.0 using a 1 cm Suprasil quartz cuvette (Starna Scientific Ltd., UK) using a Biochrom Libra 50 UV/Visible spectrometer (Biochrom Ltd., UK). IMP-1 concentrations were determined at A_260_ using Beer-Lambert with a calculated nearest neighbour extinction co-efficient (ε260) of = 113,589,000 M ^-1^ cm ^-1^ (51).

### 2.3) Dynamic Light Scattering (DLS)

DLS was measured using the UNcle ™ instrument (Unchained Labs, USA). Sample triplicates of IMP-1 samples at 0.245 mg/mL concentration were ran in UNcle quartz capillary Uni (Unchained Labs, USA), either in 5mM sodium citrate at pH 6.4 (the SAXS running buffer) or 10mM Sodium phosphate, pH 7 (the CD buffer). The DLS experiment was ran using the Uncle ™ Client software (Version 5.03) standard sizing and polydispersity method of 10 repeats every 10 second time intervals at 20 °C. The sample data was processed and averaged using the Uncle ™ Analysis software (Version 5.03) to determine the Z-Average Diameter and polydispersity index (PDI) for each buffer condition.

### 2.4) BioSAXS

BioSAXS measurements of IMP-1 were performed at 1.96 mg/mL concentration in 5 mM Sodium Citrate buffer at pH 6.4 on the SAXS-SEC using an Agilent 1260 HPLC system and SEC was performed used a Shodex KW-405 (Showa Denko K.K., Japan) column on the B21 BioSAXS beamline at the Diamond Light Source Ltd (Oxford, UK) (35) using the automated BIOSAXS robot for sample loading at 293 K. B21 was operated at a fixed camera length of 4.014 m and an energy of 12.4 keV to collect data between 0.0031 and 0.38 Å^−1^. The data was collected on a Eiger 4M detector (Dectris AG, Switzerland). The two-dimensional area detector data was converted into one-dimensional intensity profiles by radial averaging using the DAWN software package (Diamond Light Source Ltd., UK) (52). Data analysis was performed using the ATSAS v3.0 software package (53) to analyse the SEC-SAXS data and perform buffer subtraction, determine the Guinier plot and fit the PDDF (Pair-wise distance distribution function).

### 2.5) Circular dichroism (CD)

The CD samples were buffer exchanged into 10 mM Sodium Sodium Phosphate buffer at pH 7 at 0.22 mg/mL concentration determined by Beer Lambert at A_260_. All CD spectra were recorded on a nitrogen-flushed Chirascan™ Q100 spectrophotometer (Applied Photophysics Ltd., UK) in units of millidegrees (mdeg) from a wavelength range of 320 to 180 nm using a 0.05 cm path length cell (Starna Scientific Ltd., UK) and a 1 nm bandwidth, 1 nm step, 1 second integration time with 9 repeats at 20°C. The CD data was converted to units of delta Epsilon (Δε) (M^-1^ cm^-1^) after buffer spectra subtraction.

## 3) RESULTS

### 3.1) UV spectroscopy

In this paper we have used UV spectroscopy, DLS, HPLC-SEC BioSAXS and CD to characterise the purity, size and shape of the IMP-1 RNA molecule (IMP-1). On a biological scale, the IMP-1 RNA molecule is large, with a molecular weight of 3.72 MDa being composed of 11,551 base pairs. The absorbance at 260 nm (A_260_) of RNA and A_260:280_ ratio have classically between used to determine the concentration and purity of the molecule (51). The extinction co-efficient (ε260) was calculated for IMP-1 using the nearest neighbour (nn) and base composition (bc) method. Both methods gave similar values of 113.589 × 10 ^3^ M^−1^ cm^−1^ and 115.4038 × 10 ^6^, respectively. The A_260_ UV spectroscopy assay of IMP-1 (Figure 1) determined a triplicate averaged A_260_ = 1.201 +/- 0.003 (1σ) in 10 mM Tris, pH 8 buffer using a 50-fold diluted IMP-1 stock. Using the nearest neighbour (nn) co-efficient extinction co-efficient (ε_260_) a concentration of 1.96 mg/mL or 527 nM was calculated for the IMP-1 RNA stock (Table 1). An A_260_:A_280_ ratio of IMP-1 was calculated as 1.201/0.551 = 2.18. Ratio values greater than 2 are indicative of RNA free from protein contaminant (32).

**Table 1:**
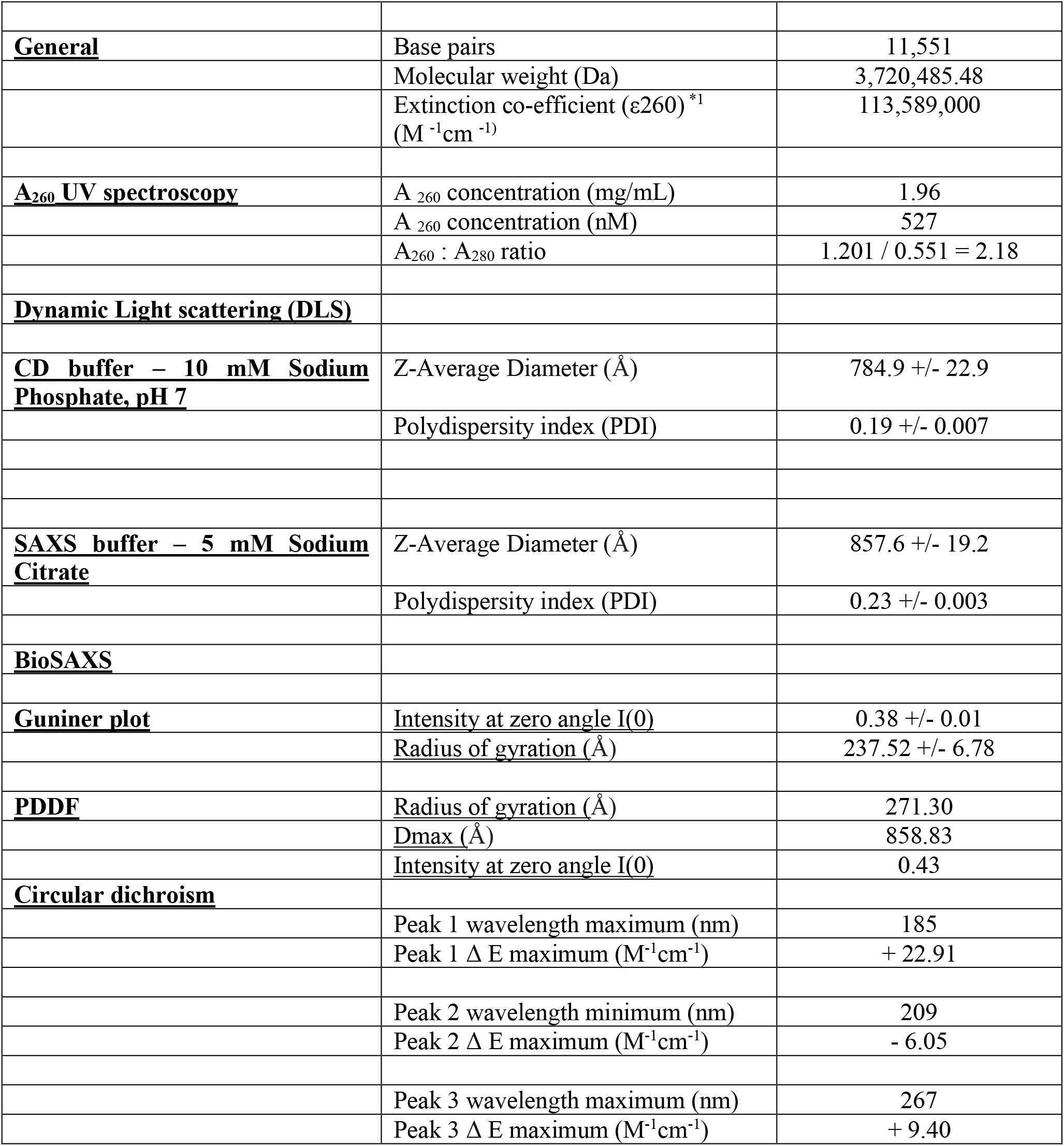
Biophysics properties of the IMP-1 RNA molecule. *1 Theoretical nearest neighbour extinction co-efficient calculated

|  |  |  |
| --- | --- | --- |
| **<u>General</u>** | Base pairs | 11,551 |
|  | Molecular weight (Da) | 3,720,485.48 |
| | Extinction co-efficient ($\epsilon_{260}$)<sup>*1</sup><br>($M^{-1}cm^{-1}$) | 113,589,000 |
| **<u>A<sub>260</sub> UV spectroscopy</u>** | A<sub>260</sub> concentration (mg/mL) | 1.96 |
|  | A<sub>260</sub> concentration (nM) | 527 |
|  | A<sub>260</sub> : A<sub>280</sub> ratio | 1.201 / 0.551 = 2.18 |
| **<u>Dynamic Light scattering (DLS)</u>** |  |  |
| **<u>CD buffer – 10 mM Sodium Phosphate, pH 7</u>** | Z-Average Diameter (Å) | 784.9 +/- 22.9 |
|  | Polydispersity index (PDI) | 0.19 +/- 0.007 |
| **<u>SAXS buffer – 5 mM Sodium Citrate</u>** | Z-Average Diameter (Å) | 857.6 +/- 19.2 |
|  | Polydispersity index (PDI) | 0.23 +/- 0.003 |
| **<u>BioSAXS</u>** |  |  |
| **<u>Guniner plot</u>** | <u>Intensity at zero angle I(0)</u> | 0.38 +/- 0.01 |
|  | <u>Radius of gyration (Å)</u> | 237.52 +/- 6.78 |
| **<u>PDDF</u>** | <u>Radius of gyration (Å)</u> | 271.30 |
|  | <u>Dmax (Å)</u> | 858.83 |
|  | <u>Intensity at zero angle I(0)</u> | 0.43 |
| **<u>Circular dichroism</u>** |  |  |
|  | Peak 1 wavelength maximum (nm) | 185 |
| | Peak 1 $\Delta E$ maximum ($M^{-1}cm^{-1}$) | + 22.91 |
|  | Peak 2 wavelength minimum (nm) | 209 |
| | Peak 2 $\Delta E$ maximum ($M^{-1}cm^{-1}$) | - 6.05 |
|  | Peak 3 wavelength maximum (nm) | 267 |
| | Peak 3 $\Delta E$ maximum ($M^{-1}cm^{-1}$) | + 9.40 |

**Figure 1:**
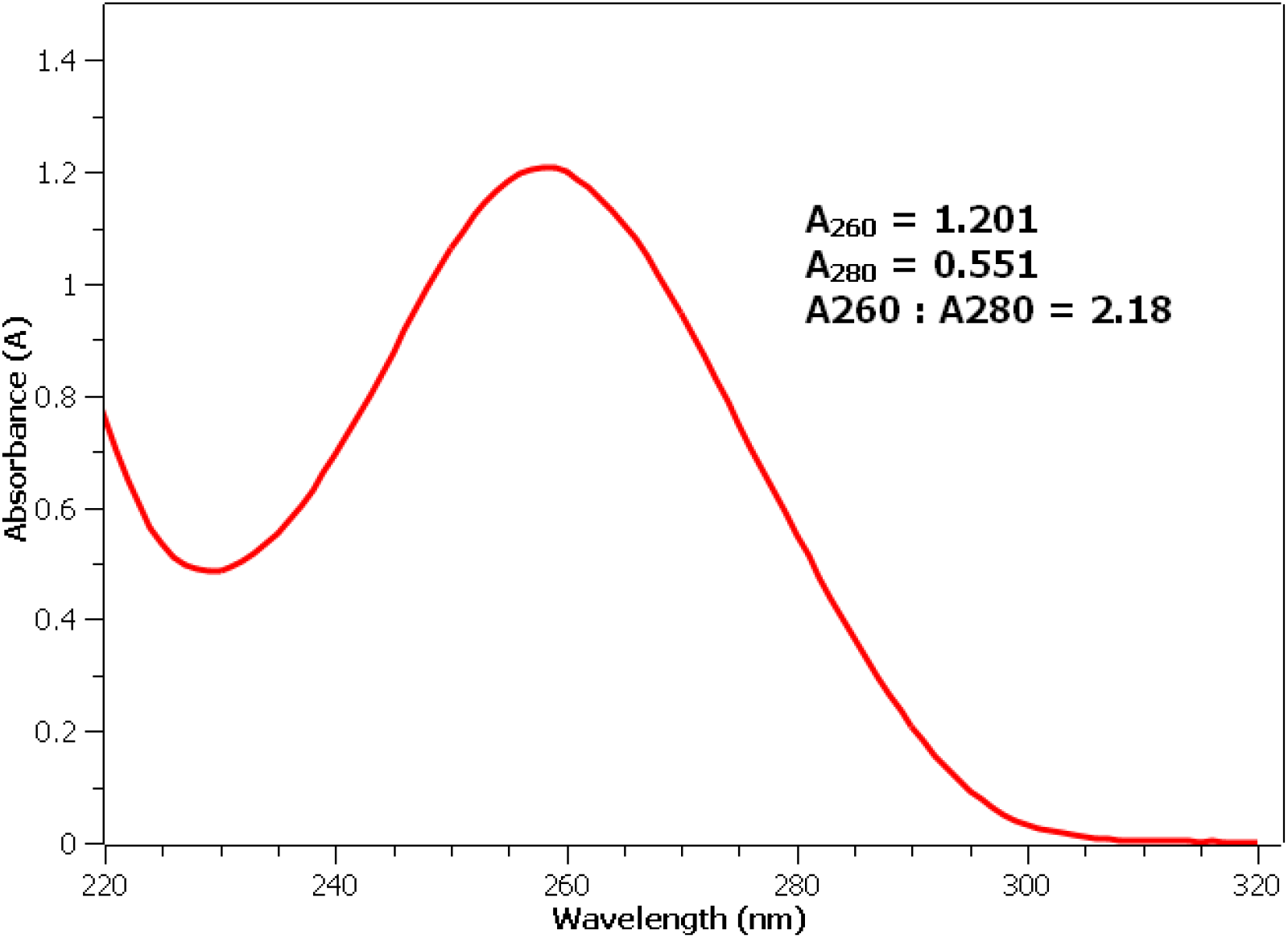
The UV spectrum of IMP-1 with the A_260_ and A_280_ values and A_260_ : A_280_ ratio given.

### 3.2) Dynamic Light Scattering

Dynamic Light Scattering (DLS) was used to determine the size of IMP-1 at 0.245 mg/mL concentration in both the SAXS running buffer (5mM Sodium Citrate, pH 6.4) and CD analysis buffer (10 mM sodium phosphate, pH 7). An intensity versus size plot of the DLS data shows a single peak for both the SAXS running buffer and CD analysis buffer data (see Figure 2). For IMP-1 in the SAXS running buffer a Z-Average hydrodynamic diameter of 857.6 +/- 19.2 Å (1σ) was determined with a corresponding polydispersity index (PDI) of 0.23 +/- 0.003 (1σ). For IMP-1 in the CD analysis a Z-Average Hydrodynamic Diameter of 784.9 Å + 22.9 Å (1σ) was determined with a corresponding polydispersity index (PDI) of 0.19 +/- 0.007 (1σ) (Table 1).

**Figure 2:**
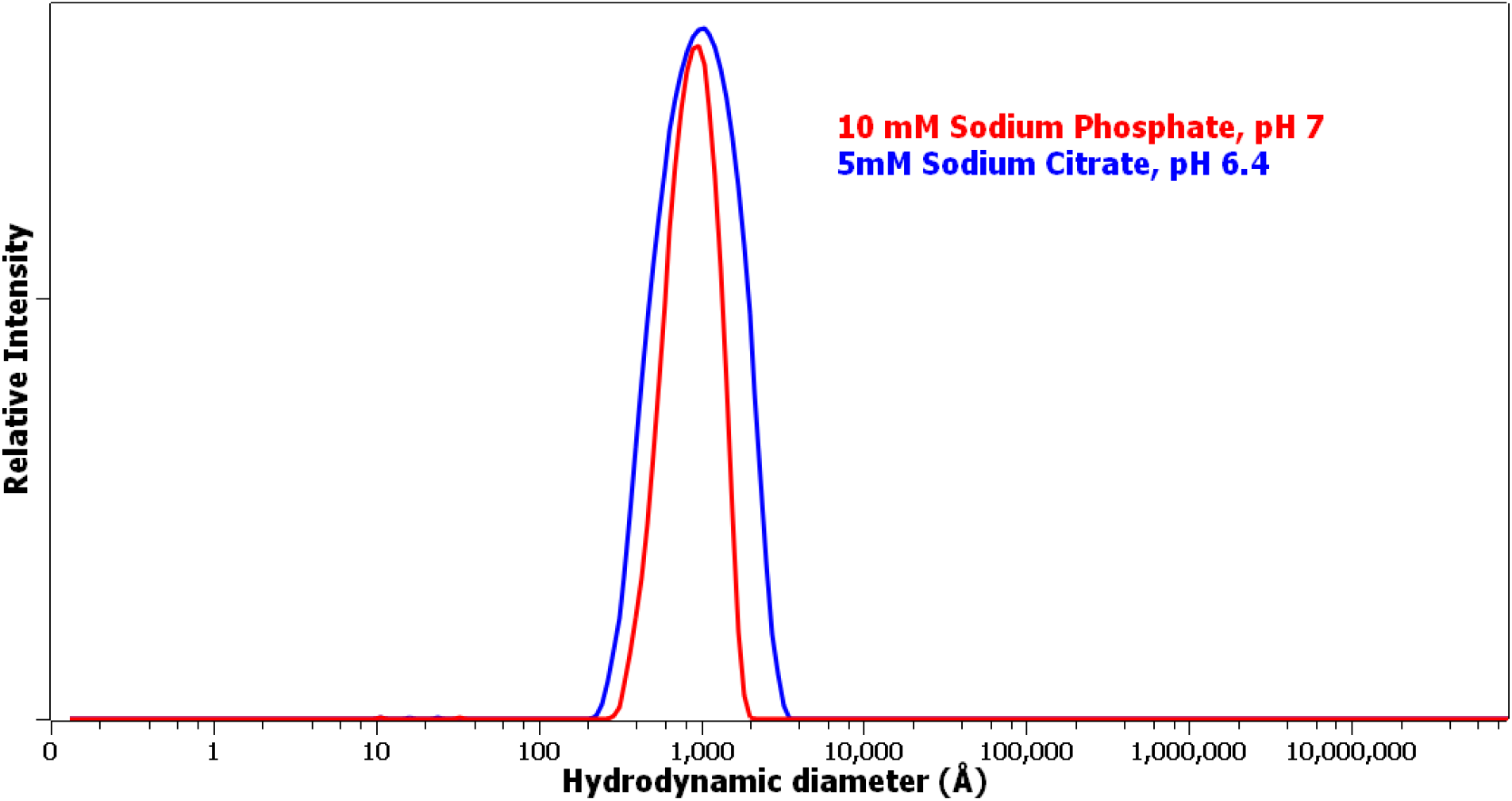
DLS spectra of the IMP-1 RNA at ∼ 0.25 mg/mL concentration in the SAXS and CD buffer.

### 3.3) BioSAXS

The BioSAXS HPLC-SEC plot of the intensity of scattering versus sample frame number resolved the IMP-1 RNA sample to a single peak using the Shodex KW-405 SEC column (Showa Denko K.K., Japan) (see Figure 3). This plot also shows the estimated radius of gyration (Rg) from a Gunier plot for each individual sample frame, with most frames within the peak giving a radius of gyration of between 200 - 250 Å (see Figure 3). From this data, sample and buffer frames were selected and an average SAXS curve for IMP-1 was generated (Figure 4). Analysis of the BioSAXS curve, using a Guinier plot gave an intensity at zero angle I(0) of 0.38 +/- 0.01 with a radius of gyration (R_g_) of 237.52 +/- 6.78 Å (Figure 4 – Inset). The IMP-1 SAXS curve was also analysed using a pair distance distribution function (PDDF) plot (see Figure 5). The PDDF plot shows a molecule which has the general shape of a solid sphere with a radius of gyration of 271.90 Å and a maximum diameter (Dmax) of 873.63 Å (Table 1). A dimensionless Kratky plot (see Figure 6) is non-parabolic and biphasic in nature suggesting the RNA molecule has both structure and flexibility.

**Figure 3:**
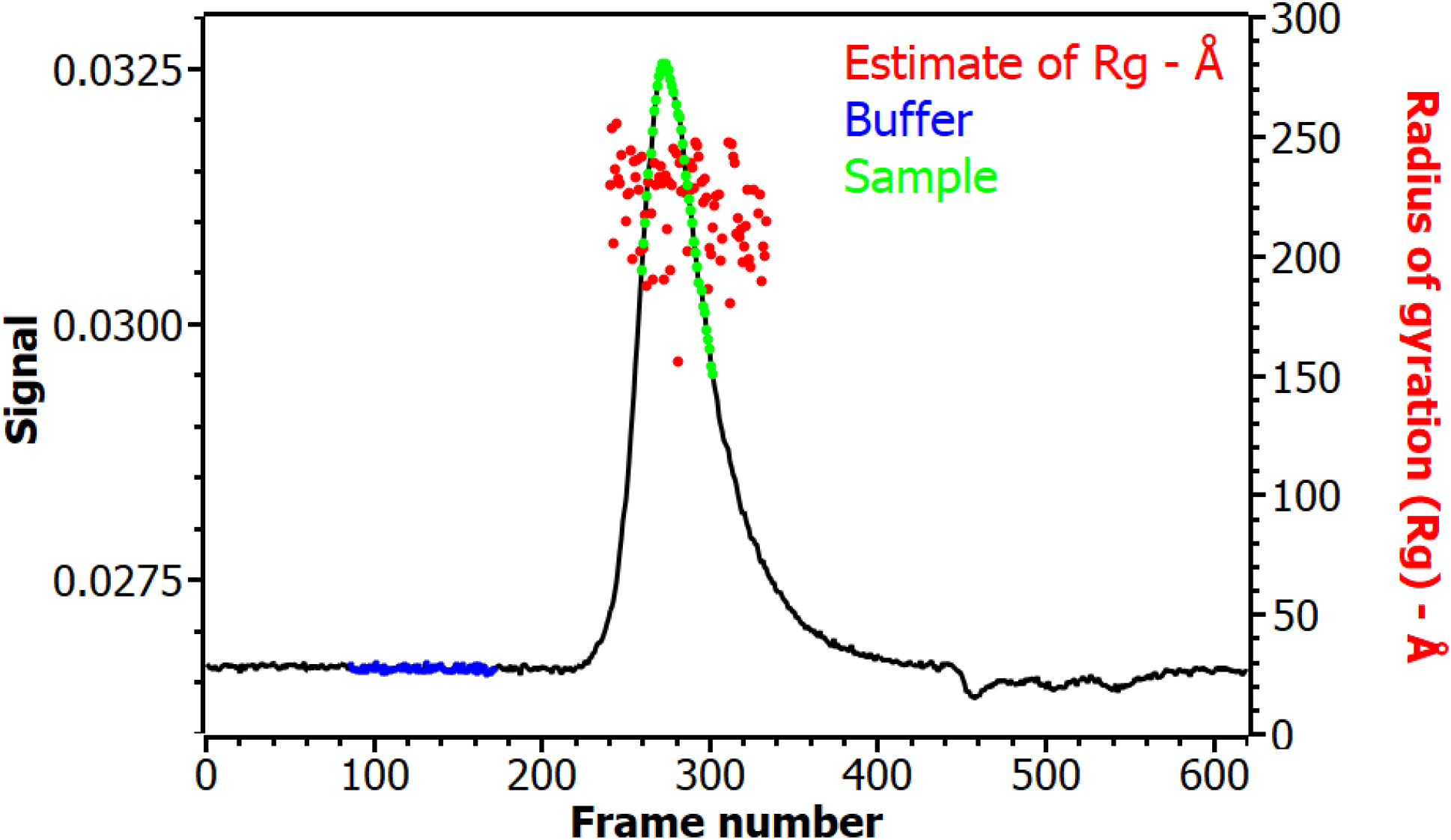
A SEC-SAXS plot of X-ray signal intensity versus frame number of IMP-1. The frames process averaged for the buffer (blue) and sample light green shown, with an estimated the radius of gyration (Rg) of each sample frame shown in red.

**Figure 4:**
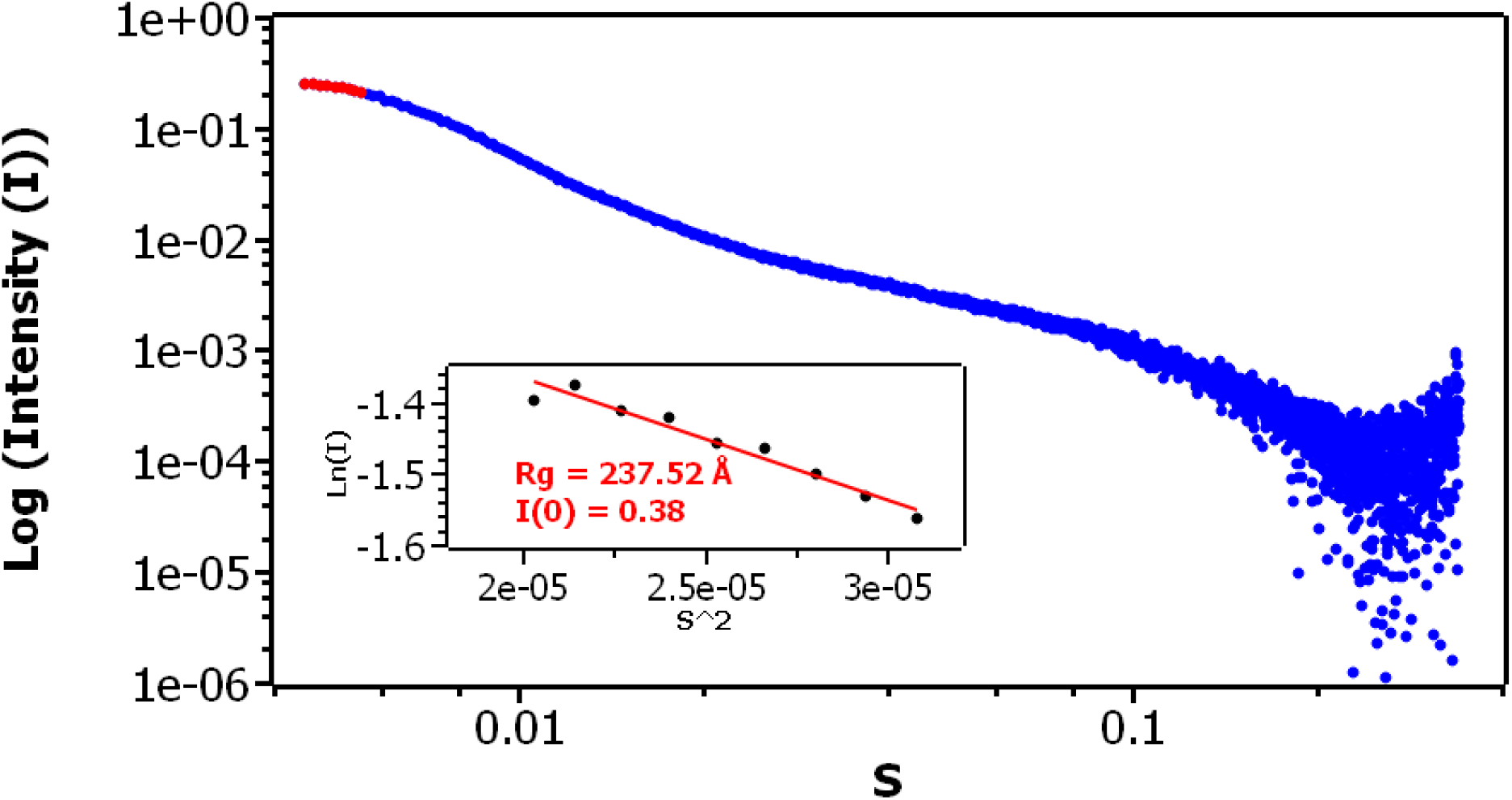
The process averaged SAXS log-log curve for IMP-1 using the buffer and sample frames shown in Figure 4, the inset shows the Guinier plot estimate of the radius of gyration (Rg) and the intensity at zero angle I(0)

**Figure 5:**
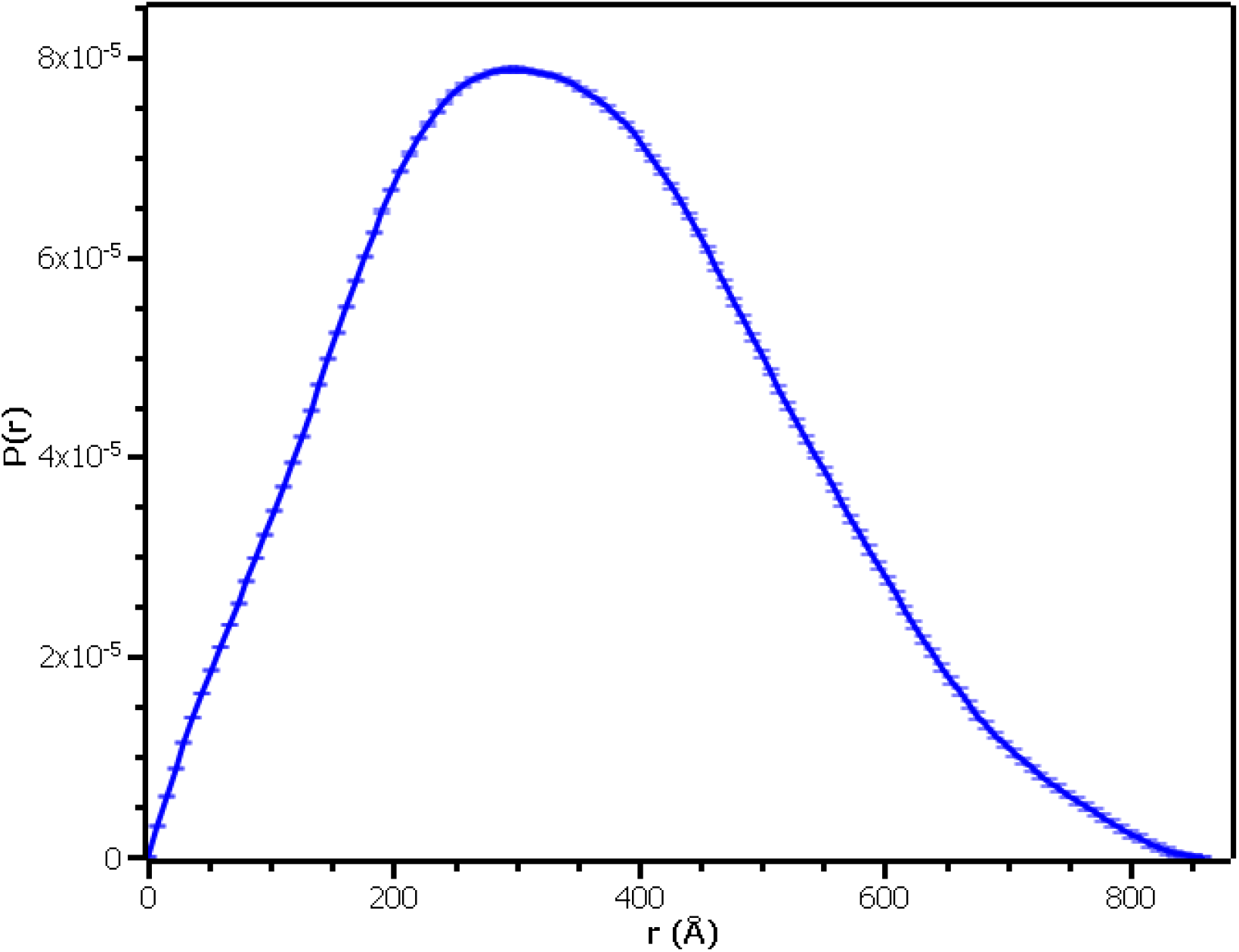
The distance (PDDF) of distribution plot the IMP-1 RNA sample taken using BioSAXS (blue – line)

**Figure 6:**
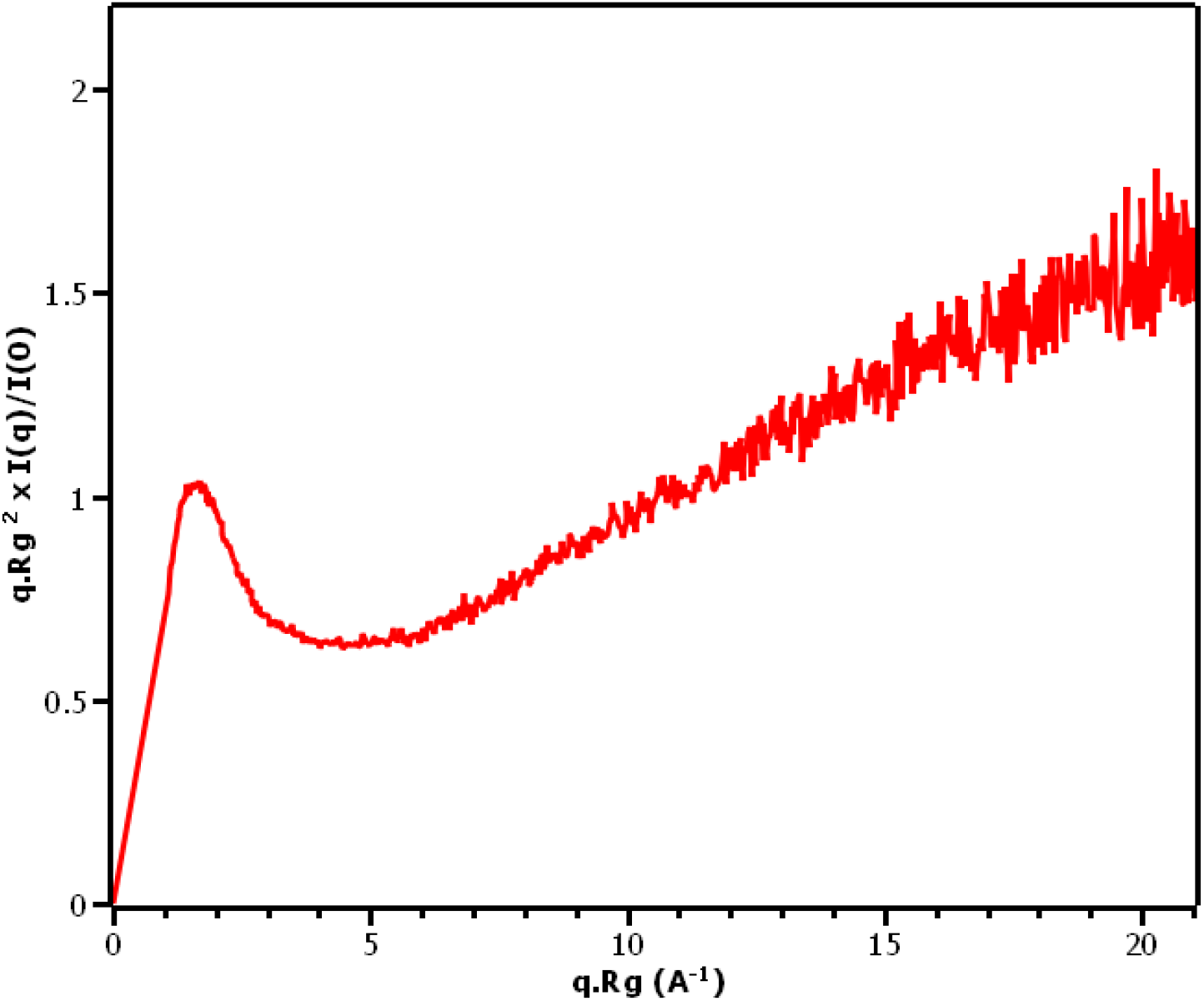
The dimensionless Kratky plot of the IMP-1 RNA sample taken using BioSAXS (blue – line)

### 3.4) Circular dichroism

The Circular dichroism spectra of IMP-1 shows structural features of the molecule (Figure 7 and Table 1). The IMP-1 CD spectra shows a positive maximum with a Δε = 22.91 M^-1^ cm^-1^ at 185 nm and corresponding negative minimum at 209 nm wavelength (Δε = - 6.05 M^-1^ cm^-1^), which is typical for right-hand A-form RNA helix (47). IMP-1 also has a positive peak at 267 nm (a Δε = - 9.40 M^-1^ cm^-1^) indicative of the sum of the base pairing and stacking interactions of the IMP-1 molecule.

**Figure 7:**
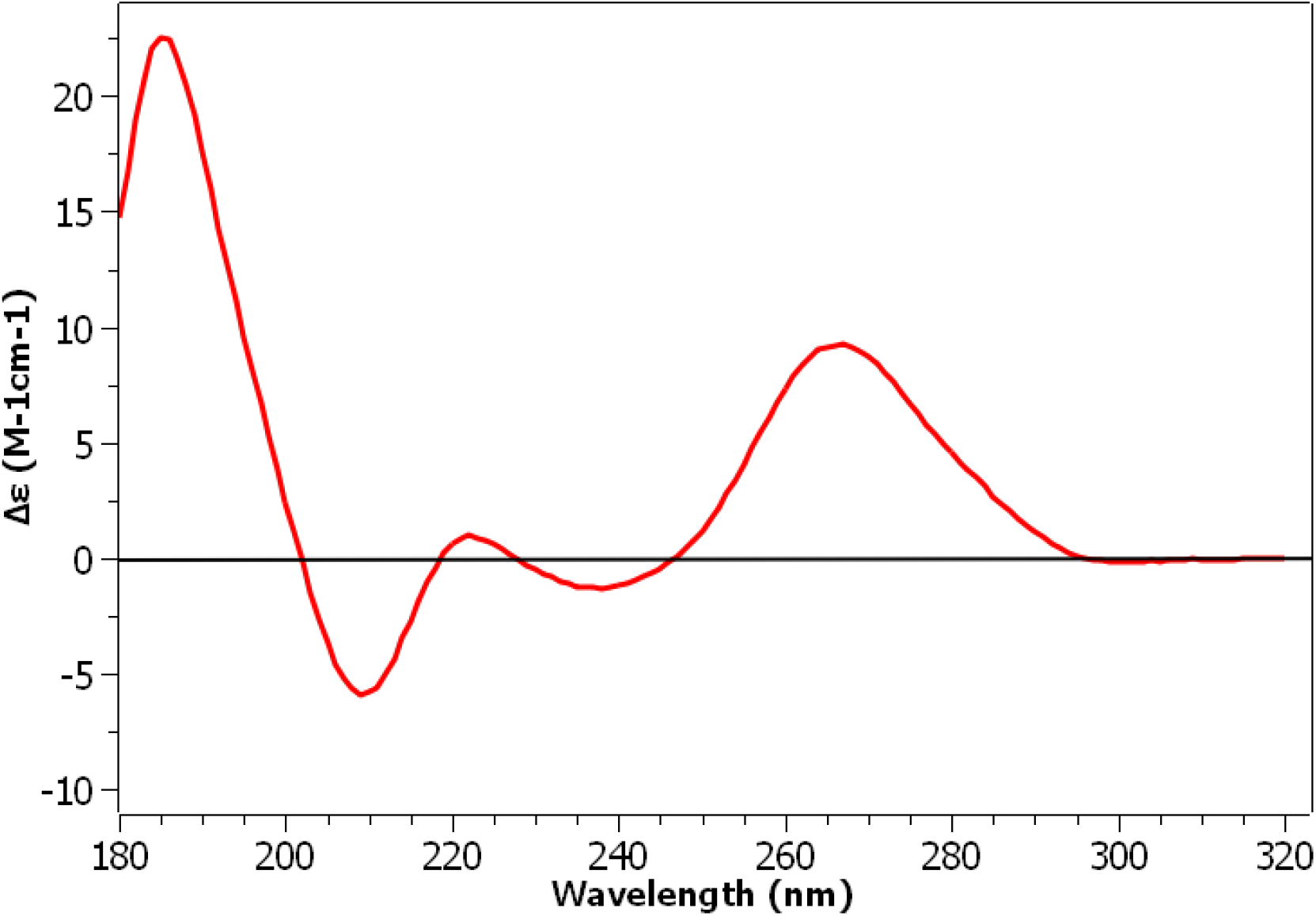
The Circular dichroism spectrum of IMP-1 in 10 mM Sodium phosphate buffer at pH 7.

## 4) DISCUSSION

RNA vaccine development understanding has increased greatly since the onset of the SARS-CoV-2 pandemic at the end of 2019. Several mRNA SARS-CoV-2 vaccines have been successfully approved or are in development. These mRNA vaccines can be split into two types, conventional and self-amplifying. Conventional mRNA vaccines, typically around 2000 bps, provide the genetic code for the host cell machinery to produce the target antigen, for SARS-CoV-2 this has typically been the surface spike protein. Self-amplifying (sa) mRNA vaccines, which has potential to be provided at significantly lower doses, encodes not only the genetic code for the target antigen, but also a viral replicase to amplify the target antigen code *in vivo*. SaRNA vaccines are therefore much larger than conventional mRNA vaccines at around 10 kbps. The development of saRNA vaccines, such as IMP-1, have proven challenging for conventional bioanalytics, due to their large size and complex structure. The development of new experimental techniques are therefore required. In this paper we have characterised several important biophysical parameters of the IMP-1 SARS-CoV-2 vaccine mRNA molecule, including its size, shape, structure and purity using UV spectroscopy, DLS and HPLC-SEC BioSAXS and circular dichroism.

UV spectroscopy is a classical method for determining the concentration and purity of IMP-1 in solution. It provides rapid determination of RNA sample concentration and the level of protein contamination (32). DLS and BioSAXS were performed to determine the size, shape and purity of the IMP-1 molecule. DLS provides the Z-Average intensity weighted mean hydrodynamic diameter (R_H_) of the total sample particles its polydispersity index (PDI) of the sample molecules. IMP-1 samples in the SAXS (5 mM Sodium Citrate, pH 4) and CD buffer (10mM Sodium phosphate, pH 7) were measured. The RNA shows small, but significant differences in Z-Average diameter and PDI between these two buffers. This suggests RNA size is dependent on buffer type, pH or ionic strength. BioSAXS gives orthogonal parameters to DLS. From the Guinier plot and PDDF a radius of gyration (Rg) of 237.52 +/- 6.78 and 271.90 Å respectively and maximum diameter (Dmax) of 873.63 Å were determined. The Dmax obtained by BioSAXS is in good agreement with the Z-Average diameter of 873.63 Å obtained by DLS. The CD spectra of IMP-1 shows structural features present in the CD buffer (10 mM Sodium phosphate buffer, pH 7) solution. Two CD peaks are of particular interest, relating to the A-form helix and base-pairing and stacking present in IMP-1. As observed in DLS, the CD of IMP-1 is highly dependent on its buffer, with the spectra obtained in water having a much smaller helical signal (data not shown). Little literature is available about the CD of RNA and none using it to characterise large mRNA molecules such as IMP-1. CD is a useful and under-appreciated technique for the characterising molecules, as it allows direct observation of the structure of RNA in solution and the effect of buffer, pH, salt, denaturants, and temperature to be observed.

The current approved mRNA vaccines from BioNTech and Moderna are conventional mRNA vaccines. Successful combined Phase I/II clinical trials of IMP-1 encapsulated in a lipid nanoparticle (LNP’s) suggest saRNA vaccines are safe to use and produces good patient responses but can be administered at significantly lower doses than conventional mRNA vaccines. The main challenge for RNA vaccines, particularly of the self-amplifying type, is the generation and administration of such a large negatively charged molecule. To counteract the negative charge, delivery of the RNA has therefore required encapsulation of the molecule in positive or ionisable lipid delivery systems, mostly commonly LNP’s. Using LNPs, this large drug product (∼ 1000 Å) can then be taken up by the cell, and the RNA release into the cytoplasm (54). Once inside the cell, the saRNA can be translated by the cell machinery. Inside the cell there are several further questions about the structure function relationship of the RNA and how this affects the translation efficiency, immunogenicity and ribonuclease protection of the molecule (5), (55), (56). Each will be important in future vaccine development.

This paper outlines the power of UV spectroscopy, DLS, BioSAXS and CD to characterise the saRNA IMP-1 molecule. This initial data suggests several areas of future development. There are general questions about the effect of pH, buffer and different cations on the size and structure of IMP-1. A screening of these variables using the techniques described is likely to be highly informative. A better understanding of the thermodynamics of the IMP-1 molecule using techniques such as UV spectroscopy, differential scanning calorimetry (DSC) and CD, would help to understand the stability and structure of the molecule at physiological relevant temperature. Techniques such as SANS, would also complement the SAXS data particularly in the formulation of LNP’s, where deuterated lipids would allow for contrast variation. Other biophysical techniques such as analytical ultracentrifuge (AUC) and electron microscopy (EM) may be useful in providing further structural information.

## ACCESSION NUMBERS

Small angle scattering data reported have been deposited with the Small Angle Scattering Biological Data Bank (SASDB) under the accession code SASDPJ3.

## AUTHOR CONTRIBUTION

DPM conceived the project. LW and CG performed the DLS experiments. DPM performed the UV and CD experiments. CP and NC performed the BioSAXS experiment. DPM processed the experimental data and prepared figures. DPM prepared the manuscript. JL, HB, RR & RJS edited the manuscript and provided comments.

## ACKNOWLEDGMENT

The authors acknowledge everyone at CPI involved in the manufacture of IMP-1 and thank Louise Taylor and Juliana Haggerty (CPI) for helpful comments and suggestions.

## REFERENCES

1. WHO Coronavirus (COVID-19) Dashboard. https://covid19.who.int

2. Krammer, F. 2020. SARS-CoV-2 vaccines in development. Nature. 586:516–527.

3. Dolgin, E. 2021. How COVID unlocked the power of RNA vaccines. Nature. 589:189–191.

4. Bloom, K., F. van den Berg, and P. Arbuthnot. 2021. Self-amplifying RNA vaccines for infectious diseases. Gene Ther. 28:117–129.

5. Blakney, A.K., S. Ip, and A.J. Geall. 2021. An Update on Self-Amplifying mRNA Vaccine Development. Vaccines. 9:97.

6. McKay, P.F., K. Hu, A.K. Blakney, K. Samnuan, J.C. Brown, R. Penn, J. Zhou, C.R. Bouton, P. Rogers, K. Polra, P.J.C. Lin, C. Barbosa, Y.K. Tam, W.S. Barclay, and R.J. Shattock. 2020. Self-amplifying RNA SARS-CoV-2 lipid nanoparticle vaccine candidate induces high neutralizing antibody titers in mice. Nat Commun. 11:3523.

7. UK Vaccine Taskforce 2020 Achievements and Future Strategy. https://assets.publishing.service.gov.uk/government/uploads/system/uploads/attachment_data/file/1027646/vtf-interim-report.pdf

8. Scheuber, A. 2020. Imperial social enterprise to accelerate low-cost COVID-19 vaccine. https://www.imperial.ac.uk/news/198053/imperial-social-enterprise-accelerate-lowcost-covid19/

9. 100 Days Mission to Respond to Future Pandemic Threats. 2021. UK Government. https://www.gov.uk/government/publications/100-days-mission-to-respond-to-future-pandemic-threats

10. Geall, A.J., C.W. Mandl, and J.B. Ulmer. 2013. RNA: The new revolution in nucleic acid vaccines. Seminars in Immunology. 25:152–159.

11. Samnuan, K., A.K. Blakney, P.F. McKay, and R.J. Shattock. 2021. Design-of-Experiments In Vitro Transcription Yield Optimization of Self-Amplifying RNA. Molecular Biology.

12. Walker, S.E., and J. Lorsch. 2013. RNA Purification – Precipitation Methods. In: Methods in Enzymology. Elsevier. pp. 337–343.

13. Rosa, S.S., D.M.F. Prazeres, A.M. Azevedo, and M.P.C. Marques. 2021. mRNA vaccines manufacturing: Challenges and bottlenecks. Vaccine. 39:2190–2200.

14. Blakney, A.K., P.F. McKay, B.I. Yus, Y. Aldon, and R.J. Shattock. 2019. Inside out: optimization of lipid nanoparticle formulations for exterior complexation and in vivo delivery of saRNA. Gene Ther. 26:363–372.

15. BioNTech COVID-19 mRNA vaccine (nucleoside-modified) EMA Procedure No. EMEA/H/C/005735/0000 Assessment report. 2020.https://www.ema.europa.eu/en/documents/assessment-report/comirnaty-epar-public-assessment-report_en.pdf

16. Demelenne, A., A.-C. Servais, J. Crommen, and M. Fillet. 2021. Analytical techniques currently used in the pharmaceutical industry for the quality control of RNA-based therapeutics and ongoing developments. Journal of Chromatography A. 1651:462283.

17. 2004. Molecular biology of the gene - Chapter 6 : The Structures of DNA and RNA. 5th ed. San Francisco: Pearson/Benjamin Cummings.

18. Varani, G., and W.H. McClain. 2000. The G·U wobble base pair: A fundamental building block of RNA structure crucial to RNA function in diverse biological systems. EMBO Rep. 1:18–23.

19. Butcher, S.E., and A.M. Pyle. 2011. The Molecular Interactions That Stabilize RNA Tertiary Structure: RNA Motifs, Patterns, and Networks. Acc. Chem. Res. 44:1302–1311.

20. Barnwal, R.P., F. Yang, and G. Varani. 2017. Applications of NMR to structure determination of RNAs large and small. Archives of Biochemistry and Biophysics. 628:42–56.

21. Garmann, R.F., A. Gopal, S.S. Athavale, C.M. Knobler, W.M. Gelbart, and S.C. Harvey. 2015. Visualizing the global secondary structure of a viral RNA genome with cryo-electron microscopy. RNA. 21:877–886.

22. Fang, X., J.R. Stagno, Y.R. Bhandari, X. Zuo, and Y.-X. Wang. 2015. Small-angle X-ray scattering: a bridge between RNA secondary structures and three-dimensional topological structures. Current Opinion in Structural Biology. 30:147–160.

23. RCSB Protein data bank: PDB Statistics: RNA-only Structures Released Per Year. https://www.rcsb.org/stats/growth/growth-rna

24. Strobel, E.J., A.M. Yu, and J.B. Lucks. 2018. High-throughput determination of RNA structures. Nat Rev Genet. 19:615–634.

25. Li, B., Y. Cao, E. Westhof, and Z. Miao. 2020. Advances in RNA 3D Structure Modeling Using Experimental Data. Front. Genet. 11:574485.

26. Ponce-Salvatierra, A., Astha, K. Merdas, C. Nithin, P. Ghosh, S. Mukherjee, and J.M. Bujnicki. 2019. Computational modeling of RNA 3D structure based on experimental data. Bioscience Reports. 39:BSR20180430.

27. Cruz-León, S., and N. Schwierz. 2020. Hofmeister Series for Metal-Cation–RNA Interactions: The Interplay of Binding Affinity and Exchange Kinetics. Langmuir. 36:5979–5989.

28. Borodavka, A., S.W. Singaram, P.G. Stockley, W.M. Gelbart, A. Ben-Shaul, and R. Tuma. 2016. Sizes of Long RNA Molecules Are Determined by the Branching Patterns of Their Secondary Structures. Biophysical Journal. 111:2077–2085.

29. Fischer, N.M., M.D. Polêto, J. Steuer, and D. van der Spoel. 2018. Influence of Na+ and Mg2+ ions on RNA structures studied with molecular dynamics simulations. Nucleic Acids Research. 46:4872–4882.

30. Tan, Z.-J., and S.-J. Chen. 2006. Nucleic Acid Helix Stability: Effects of Salt Concentration, Cation Valence and Size, and Chain Length. Biophysical Journal. 90:1175–1190.

31. Tan, Z.-J., and S.-J. Chen. 2011. Salt Contribution to RNA Tertiary Structure Folding Stability. Biophysical Journal. 101:176–187.

32. Wilfinger, W.W., K. Mackey, and P. Chomczynski. 1997. Effect of pH and Ionic Strength on the Spectrophotometric Assessment of Nucleic Acid Purity. BioTechniques. 22:474–481.

33. Stetefeld, J., S.A. McKenna, and T.R. Patel. 2016. Dynamic light scattering: a practical guide and applications in biomedical sciences. Biophys Rev. 8:409–427.

34. Svergun, D.I., M.H.J. Koch, P.A. Timmins, and R.P. May. 2013. Small angle x-ray and neutron scattering from solutions of biological macromolecules. First Edition. Oxford: Oxford University Press.

35. Cowieson, N.P., C.J.C. Edwards-Gayle, K. Inoue, N.S. Khunti, J. Doutch, E. Williams, S. Daniels, G. Preece, N.A. Krumpa, J.P. Sutter, M.D. Tully, N.J. Terrill, and R.P. Rambo. 2020. Beamline B21: high-throughput small-angle X-ray scattering at Diamond Light Source. J Synchrotron Rad. 27:1438–1446.

36. Jacques, D.A., and J. Trewhella. 2010. Small-angle scattering for structural biology-Expanding the frontier while avoiding the pitfalls: Small-Angle Scattering for Structural Biology. Protein Science. 19:642–657.

37. Rambo, R.P. 2017. Considerations for Sample Preparation Using Size-Exclusion Chromatography for Home and Synchrotron Sources. In: Chaudhuri B, I. Muñoz, S Qian, VS Urban, editors. Biological Small Angle Scattering: Techniques, Strategies and Tips. Singapore: Springer Singapore. pp. 31–45.

38. Guinier, A. 1939. La diffraction des rayons X aux très petits angles : application à l’étude de phénomènes ultramicroscopiques. Ann. Phys. 11:161–237.

39. Ochbaum, G., and R. Bitton. 2018. Using small-angle X-ray scattering (SAXS) to study the structure of self-assembling biomaterials. In: Self-assembling Biomaterials. Elsevier. pp. 291–304.

40. Putnam, C.D., M. Hammel, G.L. Hura, and J.A. Tainer. 2007. X-ray solution scattering (SAXS) combined with crystallography and computation: defining accurate macromolecular structures, conformations and assemblies in solution. Quart. Rev. Biophys. 40:191–285.

41. Brosey, C.A., and J.A. Tainer. 2019. Evolving SAXS versatility: solution X-ray scattering for macromolecular architecture, functional landscapes, and integrative structural biology. Current Opinion in Structural Biology. 58:197–213.

42. Kelly, S.M., T.J. Jess, and N.C. Price. 2005. How to study proteins by circular dichroism. Biochimica et Biophysica Acta (BBA) - Proteins and Proteomics. 1751:119–139.

43. Fasman, G.D. 2013. Circular Dichroism and the Conformational Analysis of Biomolecules. New York, NY: Springer.

44. Daniel H. A. Corrêa 1, 2 and Carlos H. I. Ramos. 2009. The use of circular dichroism spectroscopy to study protein folding, form and function. African Journal of Biochemical Research. 3:64–173.

45. Greenfield, N.J. 2006. Using circular dichroism collected as a function of temperature to determine the thermodynamics of protein unfolding and binding interactions. Nat Protoc. 1:2527–2535.

46. 2012. Comprehensive chiroptical spectroscopy. Vol. 2: Applications in stereochemical analysis of synthetic compounds, natural products, and biomolecules. Hoboken, NJ: Wiley.

47. Arluison, V., and F. Wien. 2020. RNA spectroscopy: methods and protocols - Chapter 11 : Application of Synchrotron Radiation Circular Dichroism for RNA Structural Analysis.

48. Cech, C.L., and I. Tinoco. 1977. Circular dichroism calculations for double-stranded polynucleotides of repeating sequence. Biopolymers. 16:43–65.

49. Rizzo, V., and J.A. Schellman. 1984. Matrix-method calculation of linear and circular dichroism spectra of nucleic acids and polynucleotides. Biopolymers. 23:435–470.

50. Herbert, A. 2019. Z-DNA and Z-RNA in human disease. Commun Biol. 2:7.

51. Cavaluzzi, M.J., and P.N. Borer. 2004. Revised UV extinction coefficients for nucleoside-5′-monophosphates and unpaired DNA and RNA. Nucleic Acids Research. 32:e13–e13.

52. Basham, M., J. Filik, M.T. Wharmby, P.C.Y. Chang, B. El Kassaby, M. Gerring, J. Aishima, K. Levik, B.C.A. Pulford, I. Sikharulidze, D. Sneddon, M. Webber, S.S. Dhesi, F. Maccherozzi, O. Svensson, S. Brockhauser, G. Náray, and A.W. Ashton. 2015. Data Analysis WorkbeNch (DAWN). J Synchrotron Rad. 22:853–858.

53. Manalastas-Cantos, K., P.V. Konarev, N.R. Hajizadeh, A.G. Kikhney, M.V. Petoukhov, D.S. Molodenskiy, A. Panjkovich, H.D.T. Mertens, A. Gruzinov, C. Borges, C.M. Jeffries, D.I. Svergun, and D. Franke. 2021. ATSAS 3.0 : expanded functionality and new tools for small-angle scattering data analysis. J Appl Crystallogr. 54:343–355.

54. Blakney, A. 2021. The next generation of RNA vaccines: self-amplifying RNA. The Biochemist. 43:14–17.

55. Andrzejewska, A., M. Zawadzka, and K. Pachulska-Wieczorek. 2020. On the Way to Understanding the Interplay between the RNA Structure and Functions in Cells: A Genome-Wide Perspective. IJMS. 21:6770.

56. Mauger, D.M., B.J. Cabral, V. Presnyak, S.V. Su, D.W. Reid, B. Goodman, K. Link, N. Khatwani, J. Reynders, M.J. Moore, and I.J. McFadyen. 2019. mRNA structure regulates protein expression through changes in functional half-life. Proc Natl Acad Sci USA. 116:24075–24083.

